# Single-nucleotide conservation state annotation of the SARS-CoV-2 genome

**DOI:** 10.1101/2020.07.13.201277

**Authors:** Soo Bin Kwon, Jason Ernst

## Abstract

Given the global impact and severity of COVID-19, there is a pressing need for a better understanding of the SARS-CoV-2 genome and mutations. Multi-strain sequence alignments of coronaviruses (CoV) provide important information for interpreting the genome and its variation. We apply a comparative genomics method, ConsHMM, to the multi-strain alignments of CoV to annotate every base of the SARS-CoV-2 genome with conservation states based on sequence alignment patterns among CoV. The learned conservation states show distinct enrichment patterns for genes, protein domains, and other regions of interest. Certain states are strongly enriched or depleted of SARS-CoV-2 mutations, which can be used to predict potentially consequential mutations. We expect the conservation states to be a resource for interpreting the SARS-CoV-2 genome and mutations.

## Introduction

With the urgent need to better understand the genome and mutations of SARS-CoV-2, multi-strain sequence alignments of coronaviruses (CoV) have become available^1^ where multiple sequences of CoV are aligned against the SARS-CoV-2 reference genome. Sequence alignments provide important information on the evolutionary history of different genomic bases. Such information can be useful in interpreting mutations, as for example bases with high level of sequence constraint or accelerated evolution in certain lineages have been shown to be enriched for phenotype-associated variants^2,3^. While existing systematic annotations that quantify sequence constraint from alignments^4,5^ are informative, they are limited in the information they convey on which genomic bases align and match between the reference and each sequence in the alignment, which may be useful in interrogating mutations^6^.

As a complementary approach, ConsHMM was recently introduced to systematically annotate a given genome with conservation states that capture combinatorial and spatial patterns in multi-species sequence alignment^6^. ConsHMM specifically models whether bases from non-reference sequences align and match to the reference. ConsHMM extends ChromHMM, a widely used method that uses a multivariate hidden Markov model (HMM) to learn patterns in epigenomic data *de novo* and annotate genomes based on them^7^. Previous work applying ConsHMM to multi-species alignment of other genomes have shown that the conservation states learned by ConsHMM capture various patterns in the alignment overlooked by previous methods and are useful for interpreting DNA elements and phenotype-associated variants^6,8^.

Motivated by the current need to better understand the SARS-CoV-2 genome and strain mutations, here we apply ConsHMM to two multi-strain sequence alignments of CoV that were recently made available^1^ and learn two sets of conservation states (**Fig. 1**). The first alignment is a 44-way alignment of Sarbecoviruses, a subgenus under genus Betacoronavirus, which is part of the family of Coronavirdae^9^. This alignment consists of SARS-CoV and 42 other Sarbecoviruses that infect bats aligned to the SARS-CoV-2 genome. The second alignment consists of 56 CoV that infect various vertebrates aligned to the SARS-CoV-2 genome. The vertebrate hosts include various mammals (e.g. human, bat, pangolin, mouse) and birds. Apart from the input alignments which were generated using phylogenetic trees, ConsHMM does not explicitly use any phylogenetic information. This is fitting for annotating virus genomes such as SARS-CoV-2 since frequent recombination among viruses makes it difficult to build an accurate tree^10^.

**Figure 1.**
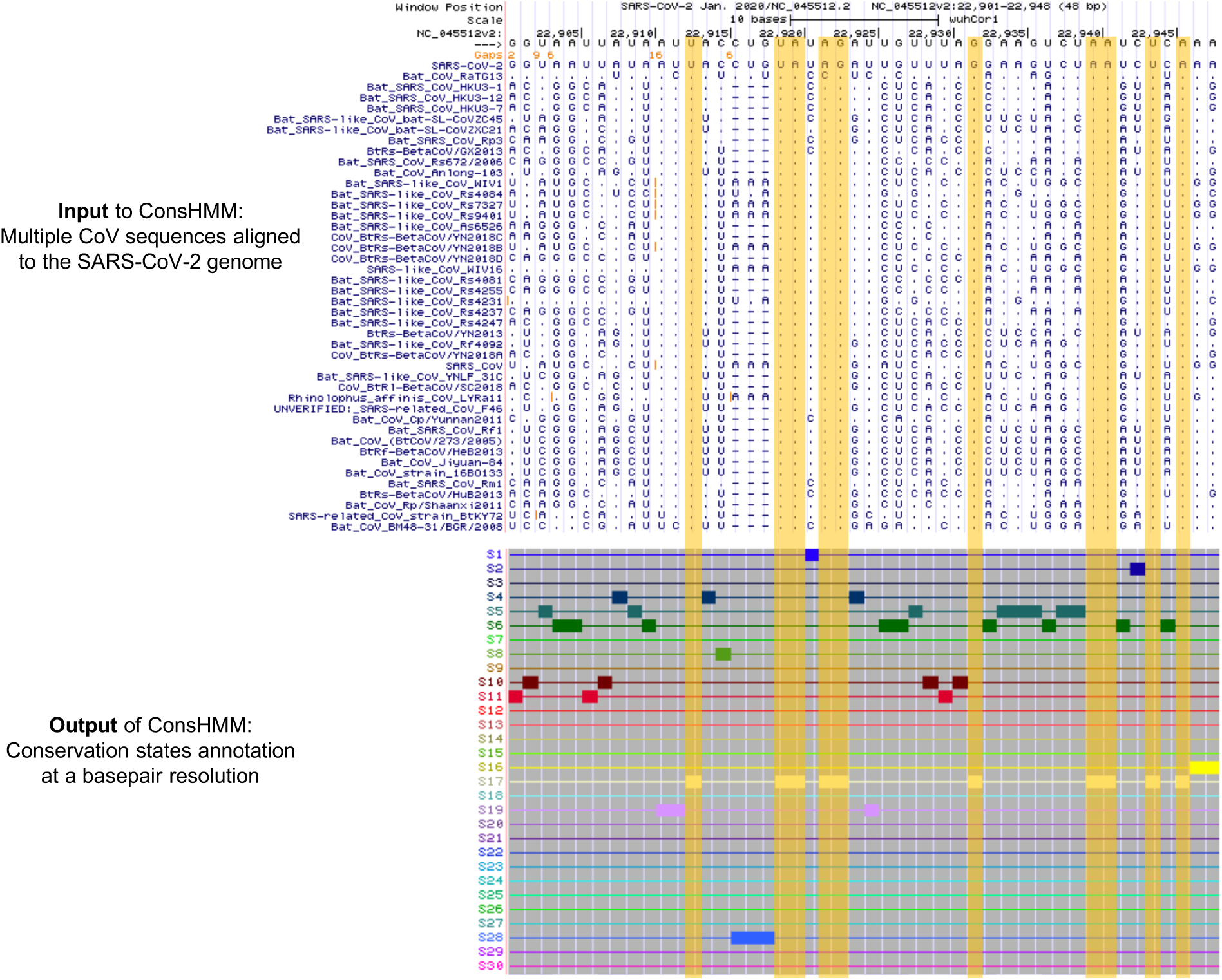
Genome browser view of ConsHMM input and output for a portion of the SARS-CoV-2 genome. Shown is an example portion of the Sarbecovirus sequence alignment input to ConsHMM and ConsHMM’s conservation state annotation of the SARS-CoV-2 genome as viewed in the UCSC Genome Browser. The top row of the alignments shows the reference sequence, the SARS-CoV-2 genome. This is followed by 43 rows corresponding to different Sarbecovirus sequences aligned against the reference, representing the 44-way Sarbecovirus sequence alignment. In each of these rows, a horizontal dash is shown if the row’s sequence has no base that aligns to the reference base shown in the top row. A dot is shown if the sequence has the same nucleotide as the reference. A specific letter is shown if for that particular base the row’s sequence has a different nucleotide than the reference. Below the alignment are 30 ConsHMM conservation states learned from the alignment. Each row corresponds to a state. To demonstrate how bases with similar alignment patterns in the input data are annotated with the same state, bases annotated with state S17 are highlighted in yellow boxes, which have most Sarbecoviruses aligning and matching to the reference with high probabilities.

Given the two sets of conservation states learned by ConsHMM from these two alignments, we annotate the SARS-CoV-2 genome with the states and analyze the states’ relationship to external annotations to understand their properties. We observe that the states capture distinct patterns in the input alignment data. Using external annotations of genes, regions of interest, and mutations observed among SARS-CoV-2 sequences, we observe that the states also have distinct enrichment patterns for various annotated regions. We generate a genome-wide track that scores each nucleotide based on state depletions and enrichments for observed mutations, which can be used to prioritize bases where mutations are more likely to be consequential. Overall, our analysis suggests that the ConsHMM conservation states highlight genomic bases with distinct evolutionary patterns in the input sequence alignments and potential biological significance. The ConsHMM conservation state annotations and the track of state depletion of mutations are resources for interpreting the SARS-CoV-2 genome and mutations.

## Result

### Annotating SARS-CoV-2 with conservation states learned from the alignment of Sarbecoviruses

First, we annotated the SARS-CoV-2 genome with 30 conservation states learned from the Sarbecovirus sequence alignment, labeled as states S1 to S30 (**Fig. 2**; **Supplementary Table 1**; **Methods**). The states capture distinct patterns of which strains align and match to the SARS-CoV-2 genome (**Fig. 2a**) and show notable enrichment patterns for external annotations of genes, proteins, and regions of interest within them (**Fig. 2b, Supplementary Table 4**). One state corresponds to bases where all strains align and match to SARS-CoV-2 with high probability and appears in the genome most frequently, covering 48% of the genome (S17). In contrast, another state corresponds to bases where only the strain closest to SARS-CoV-2, bat CoV RaTG13, aligns and matches to SARS-CoV-2 with high probability, covering 1% of the genome (S28). This state highlights bases that distinguish SARS-CoV-2 and bat CoV RaTG13 from other Sarbecoviruses. Notably, the state is highly enriched for human ACE2 binding domain (22 fold; *P*<0.0001; **Fig. 2b**), consistent with recent work suggesting that this binding domain is under strong positive selective pressure due to its critical role in host infection^11,12^. This state also annotates a region, known as the PRRA motif, that may have been inserted into the SARS-CoV-2 genome potentially resulting in increased infectiousness^13–15^. We note that this state also annotates the first five and the last seventeen bases of the genome, which may reflect technical issues with sequencing the genome ends in some strains^16^. In addition, a state corresponds to bases where all strains align to the reference with high probability but only a subset of the strains have the same nucleotide as SARS-CoV-2 with high probability (S13; **Fig. 2a**). This subset of strains includes Sarbecoviruses that are relatively distal to SARS-CoV-2 while excluding strains that are closer to SARS-CoV-2, corresponding to a deviation along a specific branch of the phylogenetic tree (**Supplementary Fig. 1**). Another state (S29) shows strong enrichment of intergenic bases (36 fold; *P*<0.0001) and gene ORF10 (59 fold; *P*<0.0001), which is consistent with recent work suggesting that ORF10 may not be a protein-coding gene based on gene expression^17^ and phylogenetic codon modeling^9^.

**Figure 2.**
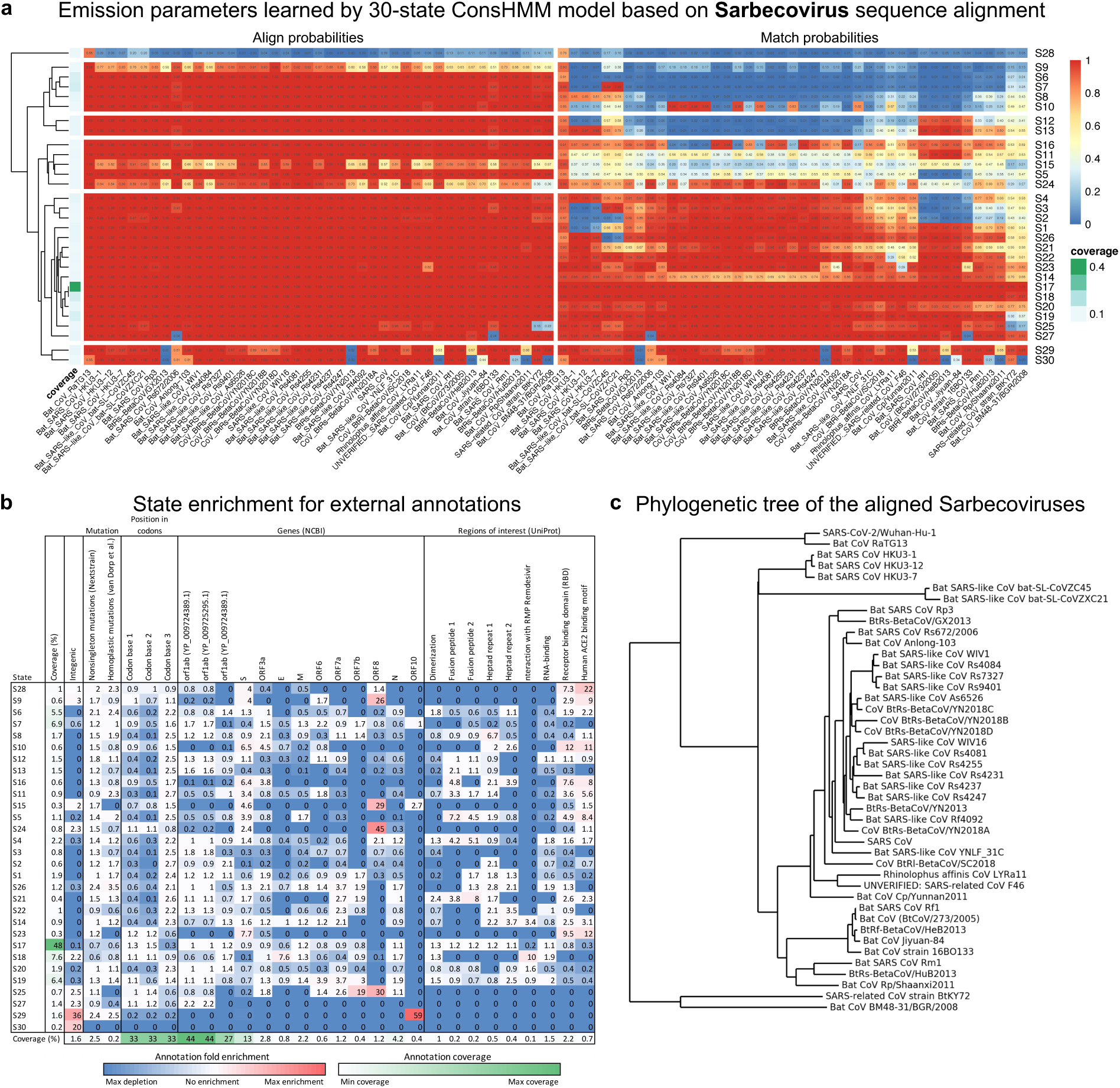
ConsHMM Conservation state learned from the Sarbecovirus alignment. **a**. State emission parameters learned by ConsHMM. The left half of the heatmap shows the probability of a base in a CoV strain aligning to the reference, which is SARS-CoV-2. The right half shows the probability of a base in a CoV strain aligning to and matching (having the same nucleotide) the reference. In both halves, each row in the heatmap corresponds to a ConsHMM conservation state with its number on the right side of the heatmap. Rows are ordered based on hierarchical clustering and optimal leaf ordering^34^. In both halves, each column corresponds to SARS-CoV or one of the 42 CoV that infect bats. Columns are ordered based on each strain’s phylogenetic distance to SARS-CoV-2, with closer strains on the left. The column on the left shows the genome-wide coverage of each state colored according to a legend labeled “coverage” on the right. **b**. State enrichment for external annotations of mutations, codons, genes, and regions of interest. The first column of the heatmap corresponds to each state’s genome coverage, and the remaining columns correspond to fold enrichments of conservation states for external annotations of intergenic regions, mutations, position within codons, NCBI gene annotations^35^, and UniProt regions of interests^21^. Each row, except the last row, corresponds to a conservation state, ordered based on the ordering shown in **a**. The last row shows the genome coverage of each annotation. Each cell corresponding to an enrichment value is colored based on its value with blue as 0 (annotation not overlapping the state), white as 1 to denote no enrichment (fold enrichment of 1), and red as the global maximum enrichment value. Each cell corresponding to a genome coverage percentage value is colored based on its value with white as 0 and green as the maximum. All annotations were accessed through UCSC Genome Browser^1^. **c**. Phylogenetic tree of the Sarbecoviruses included in the alignment. Each leaf corresponds to a Sarbecovirus strain included in the 44-way Sarbecovirus alignment. This tree was obtained from the UCSC Genome Browser^1^ and plotted using Biopython^36^. SARS-CoV-2/Wuhan-Hu-1, the reference genome of the alignment, is at the top.

### Annotating SARS-CoV-2 with conservation states learned from the alignment of Coronaviruses infecting vertebrates

In addition to the 30-state model learned from the Sarbecovirus sequence alignment, we learned another 30-state model by applying ConsHMM to the alignment of CoV from vertebrate hosts (V1∼V30; **Fig. 3**; **Supplementary Table 2**; **Methods**). The vertebrate CoV alignment consisted of a diverse set of CoV that included not only Sarbecoviruses but also CoV that are evolutionarily more diverged from SARS-CoV-2 than Sarbecoviruses. We therefore applied ConsHMM separately to the vertebrate CoV alignment, instead of combining the two alignments.

**Figure 3.**
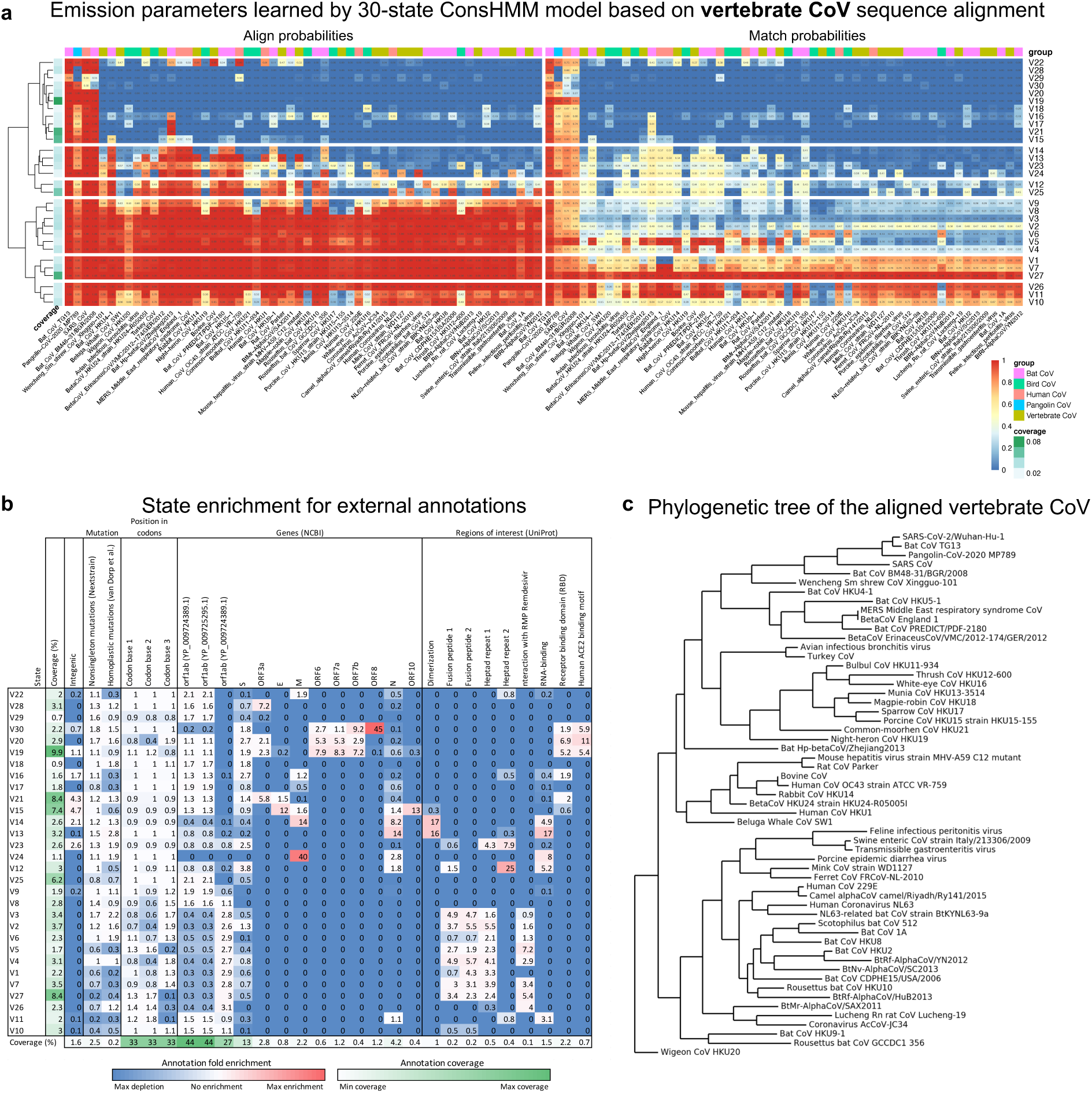
ConsHMM Conservation states learned from the vertebrate CoV alignment. **a**. State emission parameters learned by ConsHMM. The left half of the heatmap shows the probability of a base in a CoV strain aligning to the reference, which is SARS-CoV-2. The right half shows the probability of a base in a CoV strain aligning to and matching (having the same nucleotide) the reference. In both halves, each row in the heatmap corresponds to a ConsHMM conservation state with its number on the right side of the heatmap. Rows are ordered based on hierarchical clustering and optimal leaf ordering^34^. In both halves, each column corresponds to SARS-CoV or one of the 56 CoV that infect vertebrates. Columns are ordered based on each strain’s phylogenetic distance to SARS-CoV-2, with closer strains on the left. Cells in the top row above the heatmap is colored according to the color legend on the bottom right to highlight specific groups CoV with common vertebrate hosts. The column on the left shows the genome-wide coverage of each state colored according to a legend in the bottom right. **b**. State enrichment for external annotations of mutations, codons, genes, and regions of interest. The first column of the heatmap corresponds to each state’s genome coverage, and the remaining columns correspond to fold enrichments of conservation states for external annotations of intergenic regions, mutations, position within codons, NCBI gene annotations^35^, and UniProt regions of interests^21^. Each row, except the last row, corresponds to a conservation state, ordered based on the ordering shown in **a**. The last row shows the genome coverage of each annotation. Each cell corresponding to an enrichment value is colored based on its value with blue as 0 (annotation not overlapping the state), white as 1 to denote no enrichment (fold enrichment of 1), and red as the global maximum enrichment value. Each cell corresponding to a genome coverage percentage value is colored based on its value with white as 0 and green as the maximum. All annotations were accessed through UCSC Genome Browser^1^. **c**. Phylogenetic tree of the vertebrate CoV included in the alignment. Each leaf corresponds to a vertebrate CoV strain included in the vertebrate CoV. This tree was generated by pruning out SARS-CoV-2 genomes except the reference from the phylogenetic tree of the 119-way vertebrate CoV alignment obtained from the UCSC Genome Browser^1^ (**Methods**) and was plotted using Biopython^36^. SARS-CoV-2/Wuhan-Hu-1, the reference genome of the alignment, is at the top.

The resulting conservation states correspond to bases with distinct probabilities of aligning and matching to various strains of vertebrate CoV and exhibit notable enrichment patterns for previously annotated regions within genes (**Fig. 3a, Supplementary Table 4**). A state (V27) annotates bases that align and match to all 56 CoV with a genome coverage of 9%. Another state (V19) corresponds to bases that align and match specifically to four strains most closely related to SARS-CoV-2 based on phylogenetic distance, which include two bat CoV (RaTG13 and BM48-31/BGR/2008), pangolin CoV, and SARS-CoV. A state (V20) has a high align and match probabilities primarily for CoV with bat or pangolin as hosts and is enriched for the spike protein’s receptor binding domain (RBD), where a recombination event between a bat CoV and a pangolin CoV might have occurred^13^ (6.9 fold enrichment). Additionally, a state (V29) with high align and match probabilities specifically for bat CoV RaTG13 annotates the PRRA motif mentioned in the previous section, which is consistent with the possibility that the motif was recently introduced to the SARS-CoV-2 genome.

Since the input vertebrate CoV alignment includes several CoV infecting human, the states learned from this alignment can be used to investigate the varying pathogenicity among human CoV. A state (V14) corresponds to bases shared among pathogenic human CoV, including SARS-CoV-2, SARS-CoV, and Middle East respiratory syndrome-related CoV (MERS-CoV), but not shared among less pathogenic human CoV which are associated with common cold (OC43, HKU1, 229E, and NL63). Bases annotated by this state are candidates for contributing to the shared pathogenicity of SARS-CoV, SARS-CoV-2, and MERS (**Supplementary Table 5**). We compared bases annotated by this state to positions identified in previous study that located indels differentiating pathogenic CoV from common-cold-associated CoV using an alignment of 944 human CoV sequences under a supervised learning framework^18^. State V14 overlapped with two insertions identified in that study, one of which is in the nucleocapsid protein and was suggested to contribute to the virus’s pathogenicity by enhancing its nuclear localization signals^18^ (overlapping positions: 29115-29124). Moreover, using state V14 we identify additional loci potentially unique to pathogenic CoV that were not reported in the previous study (**Supplementary Table 5**). While this could be explained mostly by the different sequences included in the alignments used here and the previous study, we find among the additional loci those that are shared among all pathogenic sequences but missing in all common-cold-associated sequences according to the previous study’s human CoV alignment (**Supplementary Table 5**; **Methods**). Among such additional loci that are unique to pathogenic sequences but not previously reported is an 8-bp region (positions 28415-28423) in the nucleocapsid protein, a protein that was shown to enrich for indels specific to pathogenic CoV in the previous study. Overall, this demonstrates the conservation state annotations learned using an unsupervised approach identified additional genomic bases that may contribute to the pathogenicity of CoV infecting humans.

### Conservation states’ relationship to nonsingleton SARS-CoV-2 mutations observed in the pandemic

We next investigate how the learned conservation states relate to nonsingleton SARS-CoV-2 mutations observed in the current pandemic (**Fig. 4a**,**c**). Specifically, we analyze the state enrichment patterns for mutations observed at least twice in about four thousand SARS-CoV-2 sequences from GISAID (Global Initiative on Sharing All Influenza Data)^19^. To focus on reliable calls of mutations, we limited our analysis to nonsingleton mutations and also with genomic positions with known technical issues^16^ masked (**Methods**). In the Sarbecovirus model, as expected, states with high probabilities that all strains align and match to SARS-CoV-2 (S17, S18) are significantly depleted of mutations observed in the current pandemic (0.6-0.7 fold enrichment; *P*<0.0001) while several states (S6, S12, S19, S26, S28, S29) are significantly enriched for mutations (1.3-2.4 fold; *P<*0.001).

**Figure 4.**
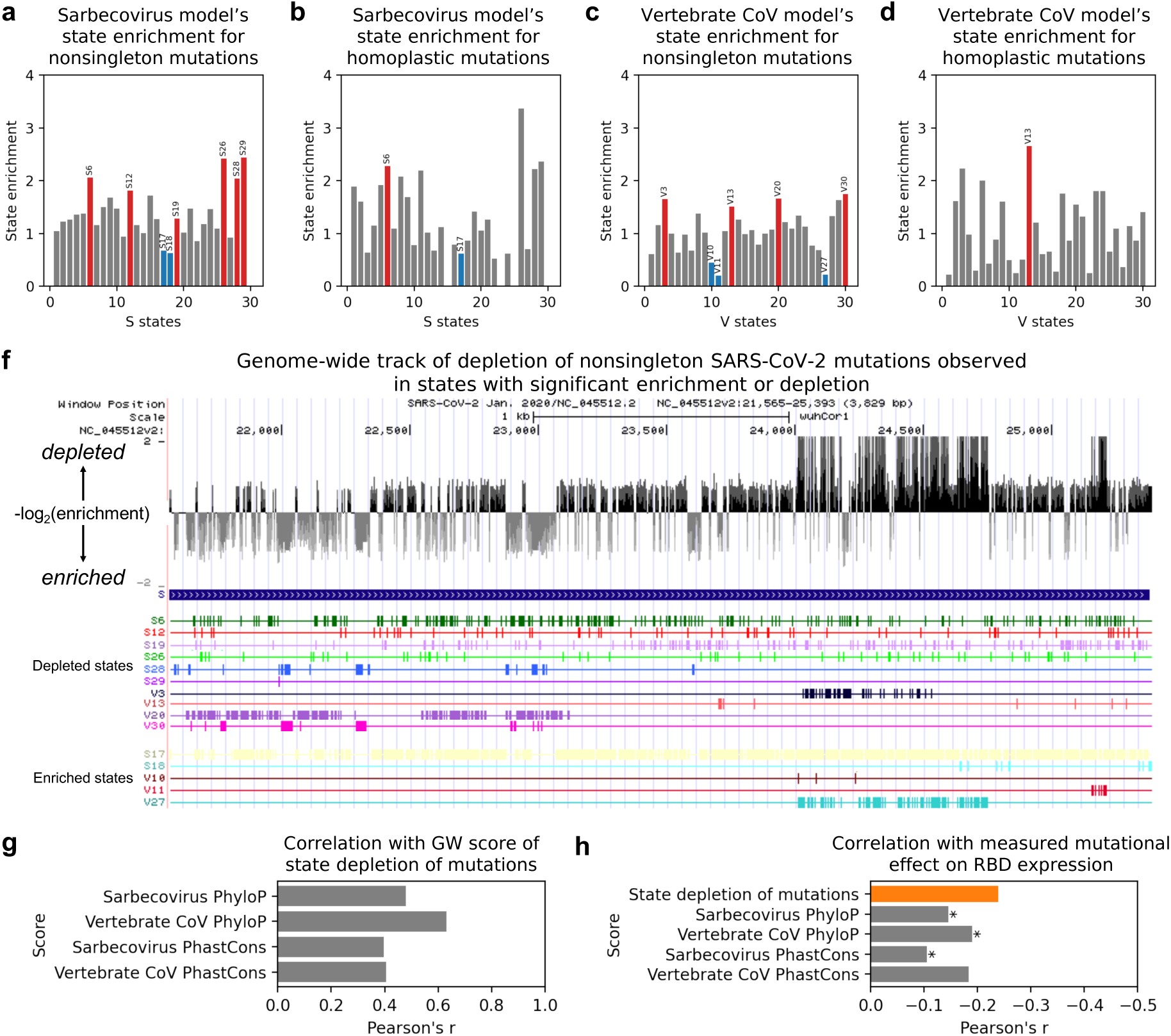
State enrichment patterns for nonsingleton mutations in the current pandemic and their relation to other annotations. **a**. Bar graph showing enrichment values of states S1-S30 learned from the Sarbecovirus sequence alignment for nonsingleton mutations (**Methods**). Red and blue bars correspond to states that enriched and depleted, respectively, with statistical significance after Bonferroni correction. Above each red or blue bar is the state ID. Grey bars correspond to states for which the enrichment was not statistically significant. Nonsingleton mutations were identified from Nextstrain mutations. **b**. Similar to **a** but showing state enrichment values for homoplastic mutations instead of nonsingleton mutations in states S1-S30. Homoplastic mutations are mutations independently and repeatedly observed in separate SARS-CoV-2 lineages and were previously stringently identified through maximum parsimony tree reconstruction and homoplasy screen using thousands of SARS-CoV-2 sequences^20^. **c**. Similar to **a** but showing state enrichment values of states V1-V30 learned from the vertebrate CoV sequence alignment instead of states S1-S30. **d**. Similar to **b** but showing state enrichment values of states V1-V30 learned from the vertebrate CoV sequence alignment instead of states S1-S30. **f**. Genome browser view of gene S with a score of depletion of nonsingleton mutations in conservation states and annotations of states from which the score is generated. Top row in black and grey vertical bars correspond to the score, which is a negative log_2_ of fold enrichment value of a state selected from either ConsHMM models that annotates a given base and is statistically significantly enriched or depleted of nonsingleton mutations at a genome-wide level (**Methods**). The following rows correspond to the states with significant enrichment or depletion. **g**. Bar graph showing correlation between our genome-wide (GW) score of state depletion of mutations shown in **f** and four sequence constraint scores listed along the y-axis. The sequence constraint scores were based on either the Sarbecovirus or vertebrate CoV sequence alignment provided to ConsHMM using either PhastCons or PhyloP as the scoring method (**Methods**). **h**. Bar graph showing correlation between measured mutational effect on RBD expression and five scores which include our state-based genome-wide score based on state depletion of mutations and the four sequence constraint scores from **g**. Correlation computed with our state-based score is shown in orange. Correlations computed with sequence constraint scores are shown in grey. Asterisk is shown next to a grey bar if its corresponding correlation was statistically significantly different than the correlation with our state-based score based on Zou’s confidence interval test^32^ with significance threshold (alpha) set to 0.0125 (0.05 divided by 4). Mutational effect on RBD expression was measured by a study that conducted a deep mutational scanning of all nonsynonymous mutations in RBD where a positive value indicates increased expression due to mutation and a negative value indicates decreased expression^24^.

The vertebrate CoV model’s conservation states exhibited some different enrichment patterns for nonsingleton SARS-CoV-2 mutations. The model learned several states that are depleted of mutations with a minimum fold enrichment of 0.2 (*P*<0.0001; V11), which is a stronger depletion than the minimum enrichment of 0.6 observed in the Sarbecovirus model. This is expected as the vertebrate CoV alignment contains a more diverse set of strains and is thus likely to capture deeper constraint than the Sarbecovirus alignment (**Fig. 3c**). Moreover, while the states significantly depleted of mutations in the Sarbecovirus model have high align and match probabilities for all strains (S17, S18), states significantly depleted of mutations in the vertebrate CoV model include not only an analogous state with high align and match probabilities for all vertebrate CoV (V27; 0.2 fold enrichment; *P*<0.0001), but also several other states that do not have high align and match probabilities for all strains (0.2-0.4 fold; P<0.0001; V10, V11). Such states have high align and match probabilities for only a subset of vertebrate CoV, which excludes strains in a specific subtree in the phylogeny of CoV, largely consisting of CoV from avian hosts (**Supplementary Fig. 2**). This indicates that bases constrained among a specific subset of vertebrate CoV, which appear to have diverged in some of the avian CoV genomes, may be as important to SARS-CoV-2 as those constrained across all vertebrate CoV. In addition, the vertebrate CoV model learns states that are enriched for mutations (1.5-1.8 fold; *P*<0.0001; V3, V13, V20, V30). The enrichment patterns for nonsingleton mutations reported here are largely consistent when we include all observed mutations or control for the nucleotide composition of each base being mutated (**Supplementary Table 3**). These patterns were also largely consistent when we control for whether each mutation is intergenic, synonymous, missense, or nonsense, indicating that the observed state enrichment patterns are not simply driven by mutation type (**Supplementary Table 3**).

To understand the state annotation’s relationship to positive selection, we next examined state enrichment patterns for homoplastic mutations (**Fig. 4b,d**). Specifically, we examined 198 stringently identified homoplastic mutations from a previous study^20^. These mutations were independently and repeatedly observed in separate SARS-CoV-2 lineages and are therefore more likely to be under positive selection than other mutations. State S6, which annotates bases with high align probability for all Sarbecoviruses but high match probability specifically for RaTG13 only, was enriched for homoplastic mutations (2.3 fold; *P*<0.001). Similarly, state V13 was significantly enriched for homoplastic mutations (2.7 fold; *P*<0.001), significantly more so than for nonsingleton mutations (1.5 fold; Binomial *P*<0.05). This state corresponds to bases that align and match to about a third of the vertebrate CoV that excludes CoV with avian host and others. The state is also enriched for the nucleocapsid protein, particularly its dimerization and RNA-binding regions which are highlighted by UniProt^21^ (14, 16, and 17 fold, respectively; *P*<0.0001).

Notably, state S17 was strongly depleted of nonsingleton mutations and homoplastic mutations (0.7 and 0.6 fold enrichment, respectively; *P*<0.0001). Interestingly, specific mutations that were previously suggested to be consequential to SARS-CoV-2 are also in this state. For example, in state S17 is a frequently observed missense mutation (position 14408) in the coding region of RNA-dependent RNA polymerase (RdRp) that was previously suggested to contribute to worsening the virus’s proofreading mechanism, making it easier for the virus to adapt and harder for its hosts to gain immunity^22^. The D614G mutation in the spike protein that was implicated to disrupt a Sarbecovirus-conserved residue^9^ and result in increased infectivty^23^ is also annotated by this state (S17). These occurrences of potentially consequential mutations in a state depleted of mutations are consistent with the notion that the state is experiencing negative selection and new mutations that do occur in the state are more likely to have stronger consequences than mutations introduced elsewhere. A similar relationship was seen with mutation type annotations, where 4% of all possible synonymous mutations were observed as nonsingleton mutations whereas only 0.3% of all possible nonsense mutations were observed as nonsigletons, reflecting their well-established difference in deleteriousness, though as noted above the conservation states show distinct enrichments for observed mutations even when conditioned on mutation type.

### Genome-wide track based on state depletion of SARS-CoV-2 mutations

We next generated a genome-wide track that reflects state depletion of mutations to highlight bases where new mutations are more likely to be consequential. The track scores each genomic base by its state’s statistically significant depletion or enrichment of nonsingleton mutations given states from both the Sarbecovirus and vertebrate CoV models, reflecting the mutation occurrence patterns among bases that likely share a common evolutionary history. To integrate the two state annotations, given two states from different ConsHMM models annotating a base of interest, we annotated the base with the state that is more depleted of nonsingleton mutations among a subset of bases that excluded the base of interest (**Methods**).

We analyzed this track based on state depletion of mutations with respect to experimentally measured mutational effect on receptor-binding domain (RBD) expression in a previous study that conducted a deep mutational scanning of RBD^24^. The study specifically measured RBD expression changes due to each possible amino acid change within RBD, where a positive value denoted increased expression and a negative value denoted decreased expression. We observed that our track based on state depletion of mutations is negatively correlated with their measured expression changes (Pearson’s *r*: -0.24, *p*<0.0001), which is consistent with our expectation that mutations at bases depleted of observed mutations in general are likely to be more deleterious than other mutations. We further compared this to four sequence constraint scores that were learned from either alignment provided to ConsHMM using PhastCons^4^ or PhyloP^5^ (**Methods**). These constraint scores were moderately correlated with our genome-wide track based on state depletion of mutations (**Fig. 4g**; Pearson’s *r*: 0.4-0.6), indicating that our track conveys distinct information. While the sequence constraint scores were also negatively correlated with the mutational effect on RBD expression, their correlations were not as strong, ranging from -0.19 to -0.11, three out of four of which were statistically significantly weaker than our track’s correlation with the mutational effect (**Fig. 4h**; *P*<0.0125; **Methods**). This suggests that our genome-wide track based on depletion of mutations could help prioritize bases to mutate when aiming to identify mutations with strong impact on the virus’s protein expression or potentially other functionalities.

## Discussion

Here we applied a comparative genomics method ConsHMM to two sequence alignments of CoV, one consisting of Sarbecoviruses that infect human and bats and the other consisting of a more diverse collection of CoV that infect various vertebrates. The conservation states learned by ConsHMM capture combinatorial and spatial patterns in the multi-strain sequence alignments. The states show associations with various other annotations not used in the model learning. The conservation state annotations are complementary to constraint scores, as they capture a more diverse set of evolutionarily patterns of bases aligning and matching, enabling one to group genomic bases by states and study each state’s functional relevance. Identifying patterns of conservation across different strains can be important potentially for understanding the relative pathogenicity of different coronaviruses and cross-immunity from prior infections^25–27^.

We showed that certain conservation states are strongly enriched or depleted of nonsingleton SARS-CoV-2 mutations. Based on this information, we generated a genome-wide track that can be used to prioritize mutations of potentially greater consequence based on evolutionary information of both the Sarbecovirus and vertebrate CoV alignments. We note that the track is generated in a transparent way directly from the fold enrichment values for nonsingleton mutations observed in the conservation states. Overall, we expect the two sets conservation state annotations along with this track based on state depletion of mutations to be resources for locating bases with distinct evolutionary patterns and analyzing mutations that are currently accumulating among SARS-CoV-2 sequences.

## Methods

### Sequence alignments

We obtained the 44-way Sarbecovirus sequence alignment from UCSC Genome Browser^1^ (**http://hgdownload.soe.ucsc.edu/goldenPath/wuhCor1/multiz44way/**). We obtained the vertebrate CoV sequence alignment by first downloading the 119-way vertebrate CoV sequence alignment from UCSC Genome Browser (**http://hgdownload.soe.ucsc.edu/goldenPath/wuhCor1/multiz119way/**) and then removing the SARS-CoV-2 sequences from the alignment, except the reference sequence, wuhCor1. This resulted in 56 CoV aligned against the reference.

### External annotations

Mutations found in SARS-CoV-2 sequences were point mutations identified by Nextstrain^28^ (accessed on Sept 7, 2020) from sequences available on GISAID^19^. For our analysis, we filtered out mutations if their ancestral alleles did not match the reference genome used by Nextstrain, MN908947.3. All the other annotations, including the annotations of genes, codons, and UniProt protein products and regions of interest, were accessed through UCSC Genome Browser (accessed on Sept 7, 2020)^1^.

### Choice of number of ConsHMM conservation states

Given the two input sequence alignments, we first learned multiple ConsHMM models from each alignment with varying numbers of states ranging from 5 to 100 with increments of 5 and then chose a number of states that is applicable to both alignments. Specifically, we aimed to find a number of states that results in states few enough to easily interpret and generalize, but specific enough to capture distinct patterns in the alignment data.

To do so, for each model, we considered whether the model’s states had sufficient coverage of the genome to avoid having states that annotate too few bases (e.g. 10 bp). We additionally considered whether the model’s states exhibited distinct emission parameters to ensure that they were different enough to capture distinct patterns in the alignment data. Lastly, we considered whether the model’s states showed distinct enrichment patterns for external annotations of genes, protein domains, and mutations in SARS-CoV-2 and showed strong predictive power for bases without mutations to ensure that the different states annotate bases with potentially different biological roles. As a result, we chose 30 as the number of conservation states for both the Sarbecovirus and vertebrate CoV ConsHMM models because the resulting states were sufficiently distinct in their emission parameters and association with external annotations and most of the states covered more than 1% of the genome.

### PhastCons and PhyloP scores

We obtained the 44-way PhastCons and PhyloP scores learned from the Sarbecovirus sequence alignment from UCSC Genome Browser (**http://hgdownload.soe.ucsc.edu/goldenPath/wuhCor1/**). We additionally used the PHAST software^29^ to learn PhastCons and PhyloP scores from the vertebrate CoV sequence alignment that we generated from the 119-way alignment as described above. To do so, we first ran ‘tree_doctor’ to prune out SARS-CoV-2 sequences except the reference from the phylogenetic tree generated for the 119-way alignment. We then followed the procedure used to generate the 44-way and 119-way scores as described on UCSC Genome Browser. Specifically, to learn the vertebrate CoV PhastCons score, we used the following arguments to run ‘phastCons’: --expected-length 45 –target-coverage 0.3 –rho 0.3. To learn the vertebrate CoV PhyloP score, we used the following arguments to run ‘phyloP’: --wig-scores –method LRT –mode CONACC.

### Masking bases

For all but one downstream analysis, we masked problematic genomic positions listed in UCSC Genome Browser track ‘Problematic Sites’ (accessed on Sept 7, 2020) as they are likely affected by sequencing errors, low coverage, contamination, homoplasy, or hypermutability^16,30,31^. The one exception was when we computed state enrichment for homoplastic mutations from a prior study. For this analysis only, we masked all problematic positions except for those described as homoplastic or highly homoplastic.

### Fold enrichment for external annotations

When computing fold enrichments for annotations of genes, positions within codons, and regions of interest, we considered whether a genomic base is annotated or not. Because multiple mutations could be observed in the same genomic base, when computing fold enrichments for mutations, we first generated all possible point mutations in the SARS-CoV-2 genome and then considered whether each of the possible mutations was observed or not. We focused on mutations observed in at least two SARS-CoV-2 sequences. For all fold enrichment values, we also conducted a two-sided binomial test to report statistical significance. We applied a Bonferroni correction by setting the significance threshold to 0.05 divided by 30, the number of states.

### Correction of state enrichment for SARS-CoV-2 mutations by nucleotide composition or mutation type

To show that the conservation state fold enrichment values for nonsingleton mutations are not simply driven by nucleotide composition or mutation type (i.e. intergenic, synonymous, missense, nonsense), we corrected state enrichment values by nucleotide composition or mutation type as follows. To control for nucleotide composition, for each nucleotide *i*, we first computed the genome-wide fraction *f*_*i*_ of observed nonsingleton mutations out of all possible mutations with nucleotide *i* as the reference base. Then for each state and for each nucleotide *i*, we multiplied the genome-wide fraction *f*_*i*_ and the number of possible mutations in the state with nucleotide *i* as the reference base. For each state, we summed up these values across the nucleotides to obtain the expected number of nonsingleton mutations based on nucleotide composition. Finally, the enrichment corrected by nucleotide composition for each state was computed as the ratio of actual and expected number of observed nonsingleton mutations.

Similarly, to control for mutation type, for each type *j* we computed the genome-wide fraction *f*_*j*_ of observed nonsingleton mutations out of all possible mutations belonging to mutation type *j*. Then for each state and for each mutation type *j*, we multiplied the genome-wide fraction *f*_*j*_ with the number of possible mutations in the state belonging to mutation type *j*. We then followed the same procedure as above.

### Identifying bases unique to pathogenic human CoV and missing in less pathogenic human CoV

We first identified bases annotated by state V14, which corresponds to high align probability for pathogenic human CoV (SARS-CoV, MERS-CoV) and low align probability for less pathogenic human CoV (OC43, HKU1, 229E, and NL63) in the vertebrate CoV sequence alignment. Among these bases, we then identified bases that appeared among all pathogenic human CoV but missing in all less pathogenic human CoV in an alignment of 944 human CoV sequences generated by a prior study. All the 944 sequences come from the seven human CoV including SARS-CoV-2^18^.

### Generating a browser track of depletion of nonsingleton SARS-CoV-2 mutations

Based on the procedure of computing state enrichment of SARS-CoV-2 mutations, we selected states from both ConsHMM models that exhibited statistically significant enrichment or depletion of nonsingleton mutations at a Binomial test p-value threshold of 0.05 after Bonferroni correction. For each base annotated with any of the selected states, we scored the base with –log_2_(*v*) where *v* is the fold enrichment value of the state annotating the base, such that stronger depletion of mutations corresponded to a higher score above 0 and stronger enrichment to a lower score below 0. If a base was annotated with two of the selected states, each from different ConsHMM models, and the two states agreed in the enrichment direction (enriched or depleted), we annotated the base with the –log_2_(*v*) from the states that had a higher absolute value of –log_2_(*v*). If a base was annotated with two of the selected states but the states disagreed in the enrichment direction, we annotated the base with score of 0. Bases not annotated by any of the selected states were assigned a score of 0 as well.

### Comparing correlation to mutational effect on RBD expression

We first computed Pearson’s *r* between the aforementioned genome-wide track based on state depletion of mutations and mutational effect on RBD expression measured by a previous study^24^. For each sequence constraint score, we computed its correlation with the measured mutation effect on RBD expression and then compared it to the correlation computed using our genome-wide track, using Zou’s confidence interval test^32^ implemented in R package cocor^33^. The four sequence constraint scores include PhyloP and PhastCons scores learned from either the Sarbecovirus or vertebrate CoV alignment. We applied a Bonferroni correction by setting the significance threshold to 0.05 divided by 4, the number of sequence constraint scores.

## Data access

ConsHMM conservation state annotation based on the Sarbecovirus and vertebrate CoV alignments are available at **https://github.com/ernstlab/ConsHMM_CoV/**. Track annotation of depletion of mutations observed in conservation states from both Sarbecovirus and vertebrate CoV ConsHMM models are available from the same URL.

## Acknowledgements

We gratefully acknowledge all those who contributed to generating and sharing their SARS-CoV-2 sequence data via the GISAID Initiative. We thank those at Nextstrain.org who made their processed mutation data publicly available. We also thank Adriana Arneson for assistance on using ConsHMM. We thank Sriram Sankararaman for comments on the manuscript. This research was supported by the UCLA David Geffen School of Medicine – Eli and Edythe Broad Center of Regenerative Medicine and Stem Cell Research Award Program and the US National Institutes of Health (DP1DA044371).

---

**Supplementary Table 1.**
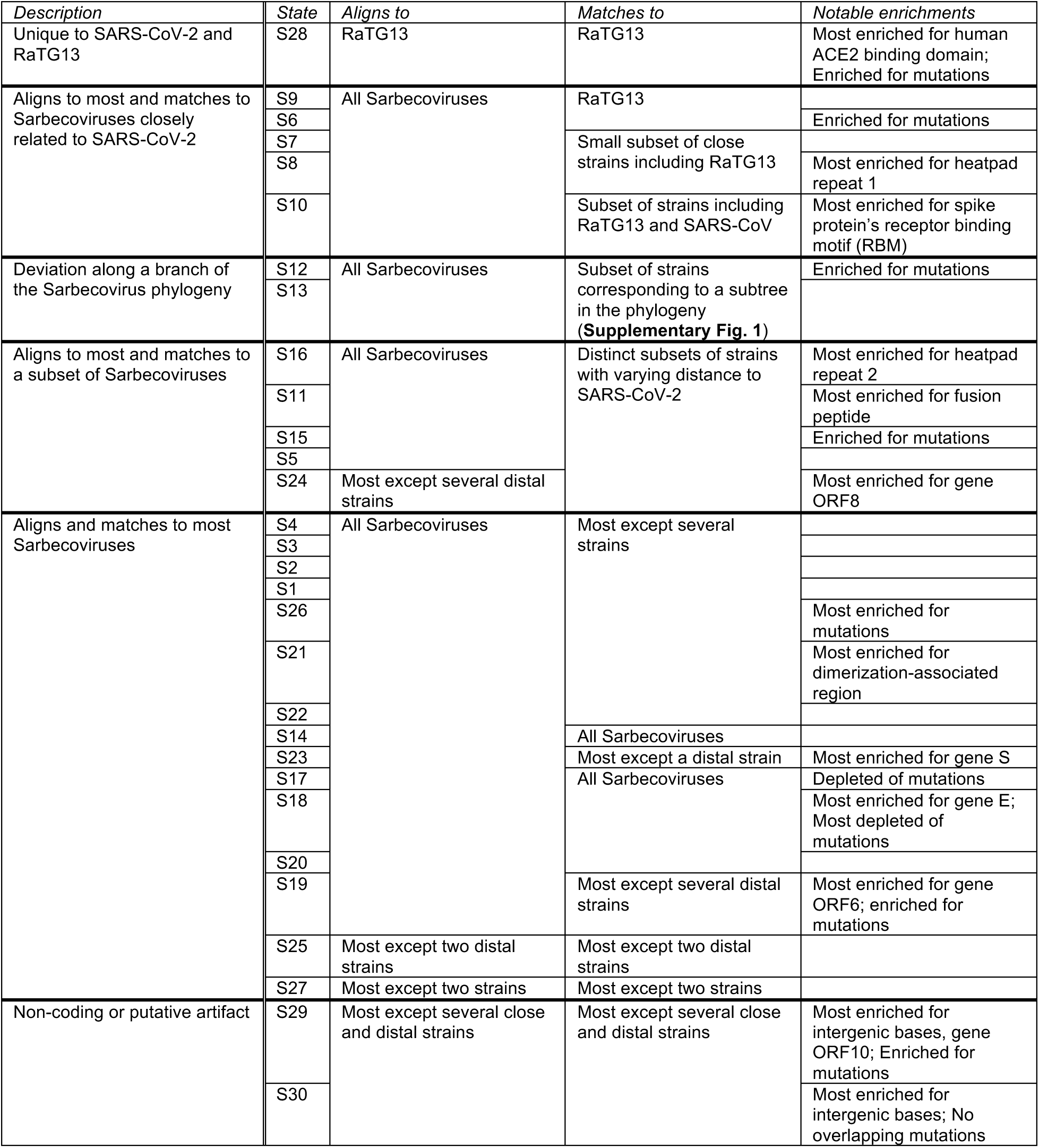
Summary of grouping, align and match probabilities, and notable enrichments of ConsHMM conservation states learned from the Sarbecovirus alignment. First column contains each group’s description, where a group consists of one or more states based on the hierarchical clustering of emission parameters as explained in **Fig. 2a**. Second column contains the state identifiers. Third and fourth columns describe the strains for which each state has align and match probabilities greater than 0.5, respectively. The last column summarizes notable enrichment of external annotations, as shown in **Fig. 2b**. RaTG13 refers to a bat CoV most closely related to SARS-CoV-2. All mutations mentioned in this table are nonsingleton mutations observed in SARS-CoV-2 sequences. All enrichment and depletion reported here have a two-sided binomial test p-value significant at a 0.05 after Bonferroni correction.

**Supplementary Table 2.**
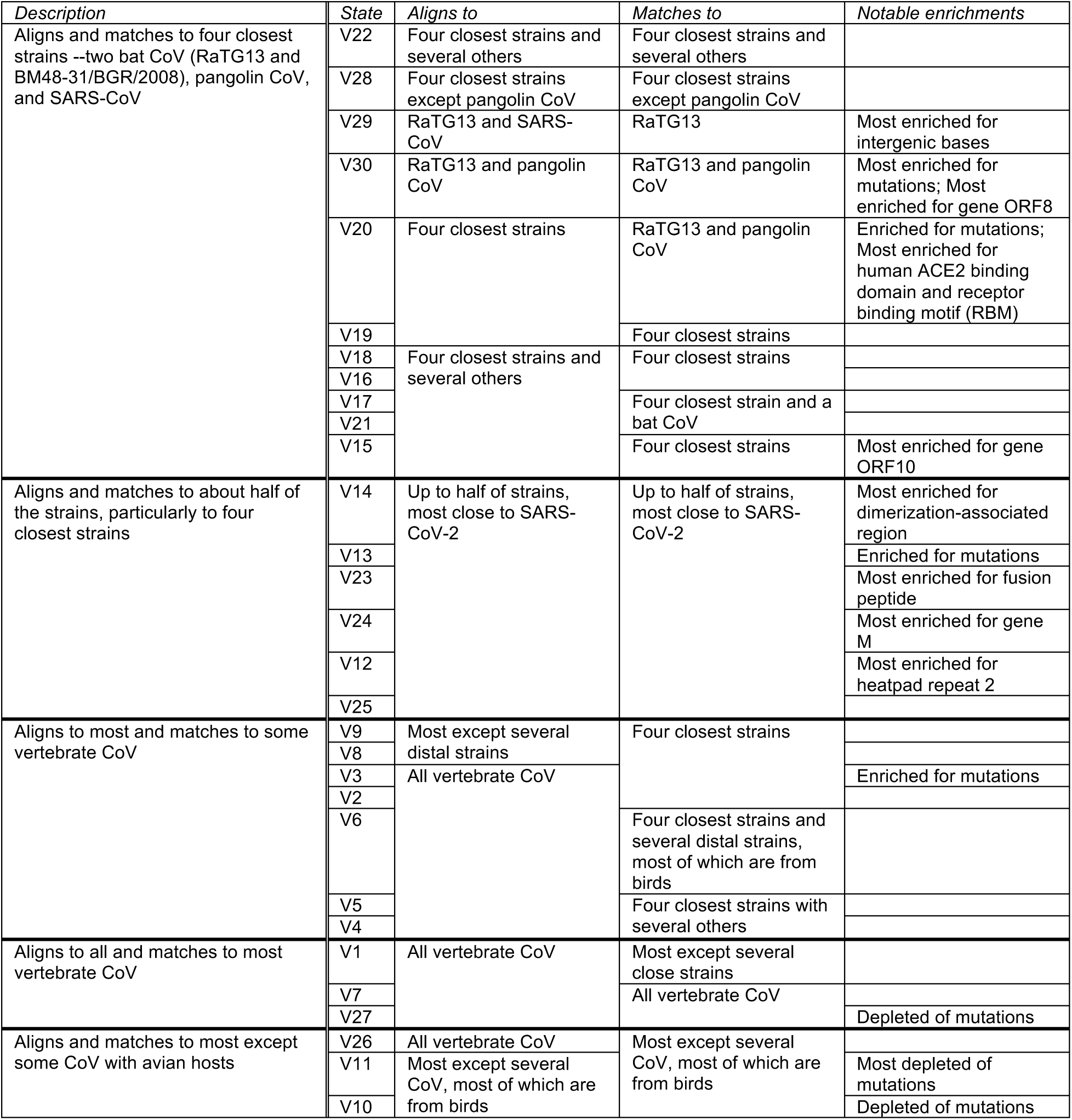
Summary of grouping, align and match probabilities, and notable enrichments of ConsHMM conservation states learned from the vertebrate CoV alignment. Similar to **Supplementary Table 1** except showing vertebrate CoV model’s states instead of Sarbecovirus model’s states.

**Supplementary Table 3.** Conservation state enrichment for SARS-CoV-2 mutations. a. Fold enrichment for SARS-CoV-2 mutations in conservation states learned from the Sarbecovirus model. Each row corresponds to a state. First column contains the state ID. State ID is shown in red if the state was significantly enriched for mutations in all six settings in which we computed enrichment which are shown in the following six columns. State ID is shown in blue if the state was significantly depleted for mutations in all settings. Otherwise, state ID is shown in black. Second column contains fold enrichment values for nonsingleton mutations currently observed in SARS-CoV-2 mutations where the enrichment is computed as the ratio between the fraction of observed mutations among possible mutations in each state and the genome-wide (GW) fraction of observed mutations among possible mutations, as done in **Fig. 2b** (**Methods**). Third column contains fold enrichment values for the same set of nonsingleton mutations except the enrichment is corrected by the nucleotide composition of the bases annotated by each state (**Methods**). Similarly, fourth column contains enrichment values for nonsingleton mutations corrected by the type (i.e. intergenic, synonymous, missense, nonsense) of the mutations annotated by each state (**Methods**). Fifth, sixth, and seventh columns are similar to second, third, and fourth columns except the enrichment values are computed based on all observed mutations instead of nonsingleton mutations. Each cell corresponding to an enrichment value is colored based on its value with blue as 0 (annotation not overlapping the state), white as 1 to denote no enrichment (fold enrichment of 1), and red as the maximum enrichment value in this table. A value is shown in bold if the associated two-sided binomial test p-value was significant at a 0.05 threshold after Bonferroni correction. **b**. Similar to **a**, except based on states learned from the vertebrate CoV model. Row order in this table do not have any correspondence to row order in **a**.

| State | Enrichment for<br>*nonsingleton* mutations |  |  | Enrichment for *all*<br>*observed* mutations |  |  |
| --- | --- | --- | --- | --- | --- | --- |
|  | Based on GW expectation | Corrected by nucleotide composition | Corrected by mutation type | Based on GW expectation | Corrected by nucleotide composition | Corrected by mutation type |
| S1 | 1.0 | 1.2 | 0.8 | 1.1 | 1.3 | 0.9 |
| S2 | 1.2 | 1.4 | 0.9 | 1.1 | 1.3 | 0.8 |
| S3 | 1.3 | 1.4 | 1.0 | 1.3 | 1.5 | 1.1 |
| S4 | 1.4 | 1.6 | 1.1 | 1.3 | 1.5 | 1.1 |
| S5 | 1.4 | 1.4 | 1.1 | 1.3 | 1.4 | 1.1 |
| S6 | 2.1 | 1.9 | 1.6 | 1.7 | 1.7 | 1.4 |
| S7 | 1.2 | 1.3 | 1.0 | 1.2 | 1.3 | 1.0 |
| S8 | 1.5 | 1.6 | 1.2 | 1.4 | 1.6 | 1.2 |
| S9 | 1.7 | 1.6 | 1.6 | 1.8 | 1.7 | 1.7 |
| S10 | 1.5 | 1.5 | 1.3 | 1.2 | 1.2 | 1.1 |
| S11 | 0.9 | 1.2 | 0.7 | 1.1 | 1.3 | 0.9 |
| S12 | 1.8 | 1.9 | 1.4 | 1.3 | 1.4 | 1.0 |
| S13 | 1.2 | 1.5 | 0.9 | 1.0 | 1.2 | 0.8 |
| S14 | 1.0 | 1.2 | 0.8 | 1.1 | 1.2 | 0.9 |
| S15 | 1.7 | 2.0 | 1.5 | 1.5 | 1.7 | 1.4 |
| S16 | 1.3 | 1.3 | 1.1 | 1.1 | 1.2 | 1.0 |
| S17 | 0.7 | 0.6 | 0.8 | 0.8 | 0.7 | 0.9 |
| S18 | 0.6 | 0.6 | 0.6 | 0.8 | 0.8 | 0.8 |
| S19 | 1.3 | 1.2 | 1.2 | 1.2 | 1.1 | 1.1 |
| S20 | 1.0 | 1.3 | 0.8 | 1.0 | 1.2 | 0.9 |
| S21 | 1.5 | 1.7 | 1.2 | 1.5 | 1.7 | 1.2 |
| S22 | 0.9 | 1.0 | 0.7 | 1.1 | 1.2 | 0.9 |
| S23 | 1.2 | 1.5 | 1.4 | 1.0 | 1.2 | 1.2 |
| S24 | 1.5 | 1.6 | 1.5 | 1.4 | 1.5 | 1.5 |
| S25 | 1.1 | 1.1 | 1.2 | 1.1 | 1.2 | 1.2 |
| S26 | 2.4 | 2.2 | 2.0 | 1.6 | 1.6 | 1.4 |
| S27 | 0.9 | 0.9 | 1.0 | 1.0 | 1.0 | 1.0 |
| S28 | 2.0 | 1.8 | 2.0 | 1.7 | 1.6 | 1.7 |
| S29 | 2.4 | 2.2 | 1.3 | 2.1 | 1.9 | 1.3 |
| S30 | 0.0 | 0.0 | 0.0 | 0.0 | 0.0 | 0.0 |

| State | Enrichment for<br>*nonsingleton* mutations |  |  | Enrichment for *all*<br>*observed* mutations |  |  |
| --- | --- | --- | --- | --- | --- | --- |
|  | Based on GW expectation | Corrected by nucleotide composition | Corrected by mutation type | Based on GW expectation | Corrected by nucleotide composition | Corrected by mutation type |
| V1 | 0.6 | 0.8 | 0.6 | 0.8 | 0.9 | 0.8 |
| V2 | 1.2 | 1.1 | 1.0 | 1.1 | 1.1 | 1.0 |
| V3 | 1.7 | 1.2 | 1.4 | 1.4 | 1.1 | 1.2 |
| V4 | 1.0 | 1.2 | 0.9 | 1.0 | 1.2 | 0.9 |
| V5 | 0.6 | 0.6 | 0.8 | 0.7 | 0.6 | 0.8 |
| V6 | 1.0 | 0.9 | 0.9 | 1.0 | 0.9 | 1.0 |
| V7 | 0.7 | 0.8 | 0.7 | 0.8 | 0.9 | 0.8 |
| V8 | 1.4 | 1.2 | 1.3 | 1.2 | 1.1 | 1.1 |
| V9 | 1.0 | 1.0 | 1.0 | 1.0 | 1.0 | 1.0 |
| V10 | 0.4 | 0.6 | 0.4 | 0.6 | 0.7 | 0.6 |
| V11 | 0.2 | 0.2 | 0.3 | 0.3 | 0.3 | 0.4 |
| V12 | 1.0 | 1.1 | 1.1 | 1.0 | 1.0 | 1.0 |
| V13 | 1.5 | 1.3 | 1.5 | 1.4 | 1.3 | 1.4 |
| V14 | 1.2 | 1.1 | 1.2 | 1.2 | 1.1 | 1.2 |
| V15 | 1.0 | 1.0 | 0.9 | 1.1 | 1.1 | 1.0 |
| V16 | 1.1 | 1.1 | 1.1 | 1.1 | 1.2 | 1.1 |
| V17 | 0.8 | 0.8 | 0.8 | 0.8 | 0.8 | 0.8 |
| V18 | 1.0 | 0.9 | 1.0 | 1.0 | 0.9 | 1.0 |
| V19 | 1.1 | 1.1 | 1.1 | 1.1 | 1.1 | 1.1 |
| V20 | 1.7 | 1.5 | 1.4 | 1.6 | 1.6 | 1.4 |
| V21 | 1.2 | 1.2 | 1.1 | 1.1 | 1.1 | 1.1 |
| V22 | 1.1 | 1.1 | 1.1 | 0.9 | 1.0 | 1.0 |
| V23 | 1.3 | 1.3 | 1.3 | 1.3 | 1.2 | 1.2 |
| V24 | 1.1 | 1.1 | 1.2 | 0.9 | 0.9 | 1.0 |
| V25 | 0.8 | 0.8 | 0.8 | 0.8 | 0.8 | 0.8 |
| V26 | 0.7 | 0.7 | 0.9 | 0.6 | 0.7 | 0.8 |
| V27 | 0.2 | 0.2 | 0.3 | 0.4 | 0.4 | 0.5 |
| V28 | 1.3 | 1.3 | 1.3 | 1.4 | 1.4 | 1.4 |
| V29 | 1.6 | 1.5 | 1.7 | 1.4 | 1.3 | 1.4 |
| V30 | 1.8 | 1.8 | 1.8 | 1.6 | 1.6 | 1.6 |

**Supplementary Table 4.**
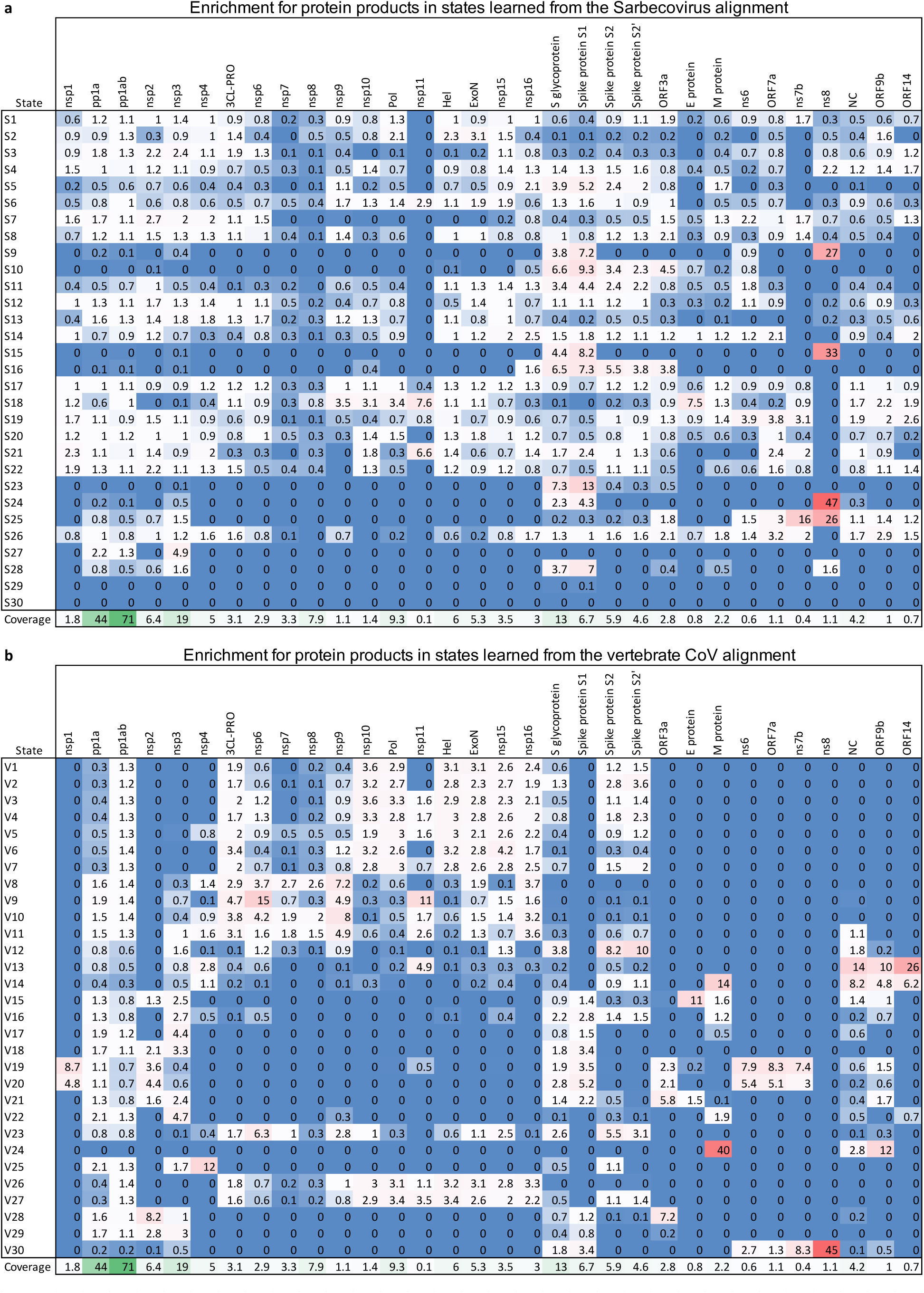
Conservation state enrichment for protein products. **a**. Fold enrichment for protein products in conservation states learned from the Sarbecovirus model. Each row corresponds to a state. First column contains the state ID. The following columns contain fold enrichment values for different protein products listed at the top of each column. Protein product coordinates and names were from UniProt Protein Product annotation^21^. Last row reports genome coverage percentage of each protein. Each cell corresponding to an enrichment value is colored based on its value with blue as 0 (annotation not overlapping the state), white as 1 to denote no enrichment (fold enrichment of 1), and red as the maximum enrichment value in this table. Each cell corresponding to a coverage percentage is colored based on its value with white as minimum and green as maximum. **b**. Similar to **a**, except based on states learned from the vertebrate CoV model.

**Supplementary Table 5.** Genomic segments unique to pathogenic human CoV and missing in less pathogenic human CoV identified by state V14. Each row corresponds to a genomic segment annotated by state V14, which corresponds to bases with high (>0.5) align probabilities for SARS-CoV and MERS-CoV and low (<0.5) align probabilities for common-cold-associated human CoV. First and second columns denote 0-based genomic coordinates (BED format). Third column shows the gene in which the genomic segments are located if it is in a gene or “non-coding” if it is not a gene. Fourth column denotes whether the base is confirmed to be unique to pathogenic human CoV and missing in less pathogenic human CoV based on an alignment of 944 human CoV sequences. Last column denotes whether the genomic segment was identified as an insertion specific to pathogenic strains in a prior study.

| start | end | gene | confirmed based<br>on human CoV | Gussow et al. |
| --- | --- | --- | --- | --- |
| 7390 | 7450 | orf1ab |  |  |
| 7807 | 7809 | orf1ab |  |  |
| 7809 | 7816 | orf1ab | TRUE |  |
| 7816 | 7825 | orf1ab |  |  |
| 7868 | 7871 | orf1ab |  |  |
| 7931 | 7933 | orf1ab |  |  |
| 8575 | 8589 | orf1ab |  |  |
| 8640 | 8647 | orf1ab |  |  |
| 8658 | 8660 | orf1ab |  |  |
| 8888 | 8892 | orf1ab |  |  |
| 8892 | 8893 | orf1ab | TRUE |  |
| 8893 | 8899 | orf1ab |  |  |
| 8963 | 8968 | orf1ab |  |  |
| 8969 | 8973 | orf1ab |  |  |
| 10237 | 10238 | orf1ab |  |  |
| 10797 | 10799 | orf1ab |  |  |
| 10869 | 10871 | orf1ab |  |  |
| 11074 | 11076 | orf1ab |  |  |
| 11370 | 11371 | orf1ab |  |  |
| 12912 | 12913 | orf1ab |  |  |
| 13328 | 13331 | orf1ab | TRUE |  |
| 16190 | 16193 | orf1ab |  |  |
| 18171 | 18174 | orf1ab |  |  |
| 18230 | 18231 | orf1ab |  |  |
| 19131 | 19134 | orf1ab |  |  |
| 19958 | 19961 | orf1ab |  |  |
| 20351 | 20353 | orf1ab |  |  |
| 20391 | 20397 | orf1ab |  |  |
| 23843 | 23844 | S |  |  |
| 23938 | 23941 | S |  |  |
| 24001 | 24002 | S |  |  |
| 24226 | 24227 | S |  |  |
| 24227 | 24228 | S | TRUE |  |
| 24228 | 24229 | S | TRUE | TRUE |
| 24775 | 24778 | S |  |  |
| 24990 | 25000 | S |  |  |
| 25322 | 25345 | S |  |  |
| 26610 | 26611 | M |  |  |
| 26874 | 26938 | M |  |  |
| 26939 | 27041 | M |  |  |
| 27043 | 27047 | M |  |  |
| 27049 | 27067 | M |  |  |
| 27078 | 27085 | M |  |  |
| 27086 | 27135 | M |  |  |
| 28396 | 28415 | N |  |  |
| 28415 | 28423 | N | TRUE |  |
| 28496 | 28500 | N |  |  |
| 28561 | 28567 | N |  |  |
| 28680 | 28686 | N |  |  |
| 28704 | 28706 | N |  |  |
| 28797 | 28809 | N |  |  |
| 28857 | 28875 | N |  |  |
| 28946 | 28966 | N |  |  |
| 29001 | 29002 | N |  |  |
| 29012 | 29014 | N |  |  |
| 29024 | 29026 | N |  |  |
| 29115 | 29116 | N |  | TRUE |
| 29116 | 29124 | N | TRUE | TRUE |
| 29218 | 29233 | N |  |  |
| 29241 | 29362 | N |  |  |
| 29374 | 29400 | N |  |  |
| 29730 | 29731 | non-coding |  |  |
| 29764 | 29771 | non-coding |  |  |
| 29784 | 29803 | non-coding |  |  |

**Supplementary Figure 1.**
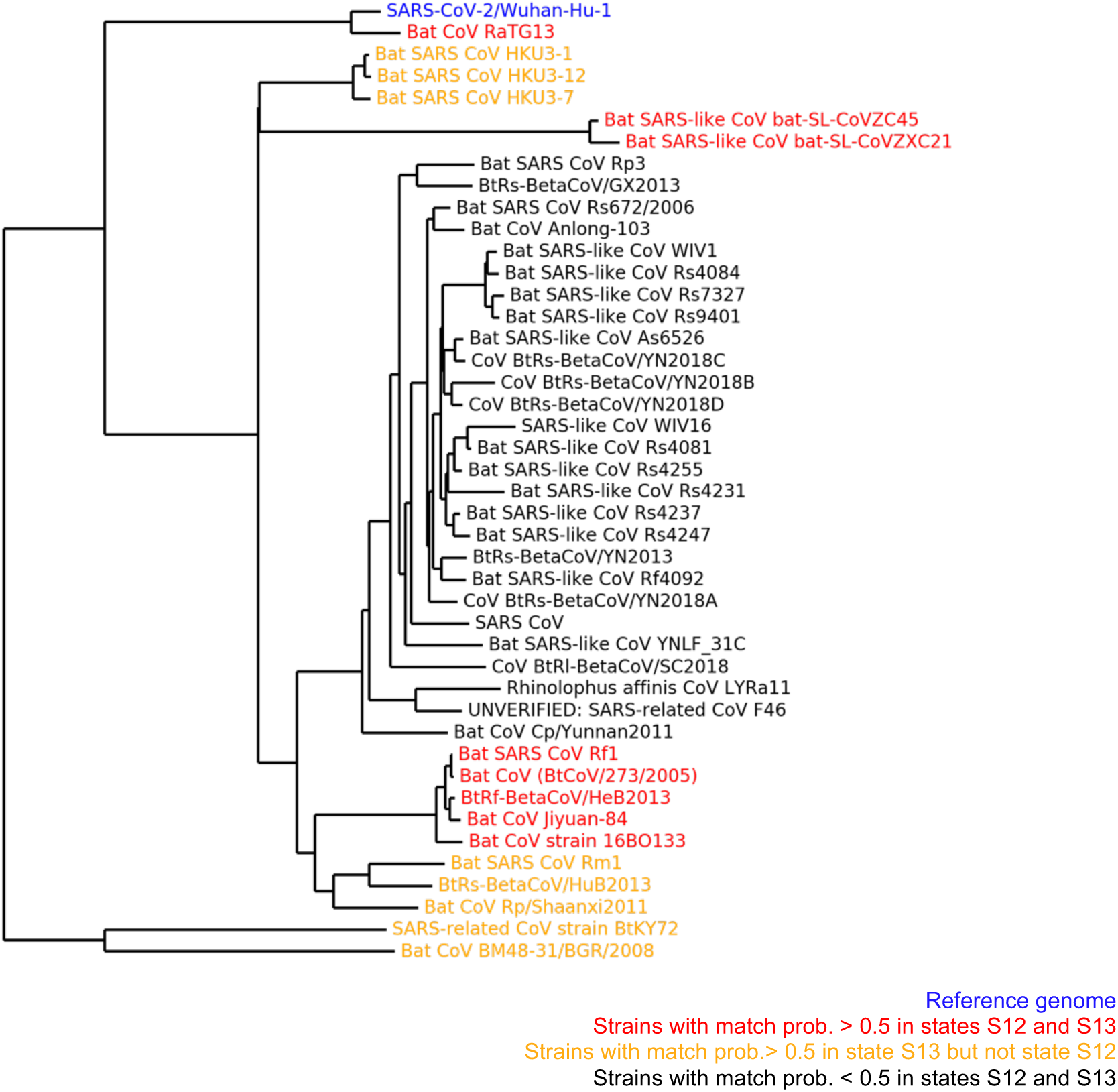
Sarbecoviruses associated with states S12 and S13 in the phylogenetic tree of the 44-way Sarbecovirus alignment. Similar to **Fig. 2c** except strains colored according to their align and match probabilities in states S12 and S13. The strain colored in blue is the reference SARS-CoV-2 strain of the alignment, SARS-CoV-2/Wuhan-Hu-1. Strains colored in black are those that have match probabilities below 0.5 for both states S12 and S13. Strains colored in red are those with match probabilities above 0.5 for both states S12 and S13. Strains colored in yellow are those with match probabilities above 0.5 for state S13 but not for state S12. All strains have high (>0.95) align probabilities for states S12 and S13. States S12 and S13 are likely to correspond to a deviation along the branch preceding all strains colored in black.

**Supplementary Figure 2.**
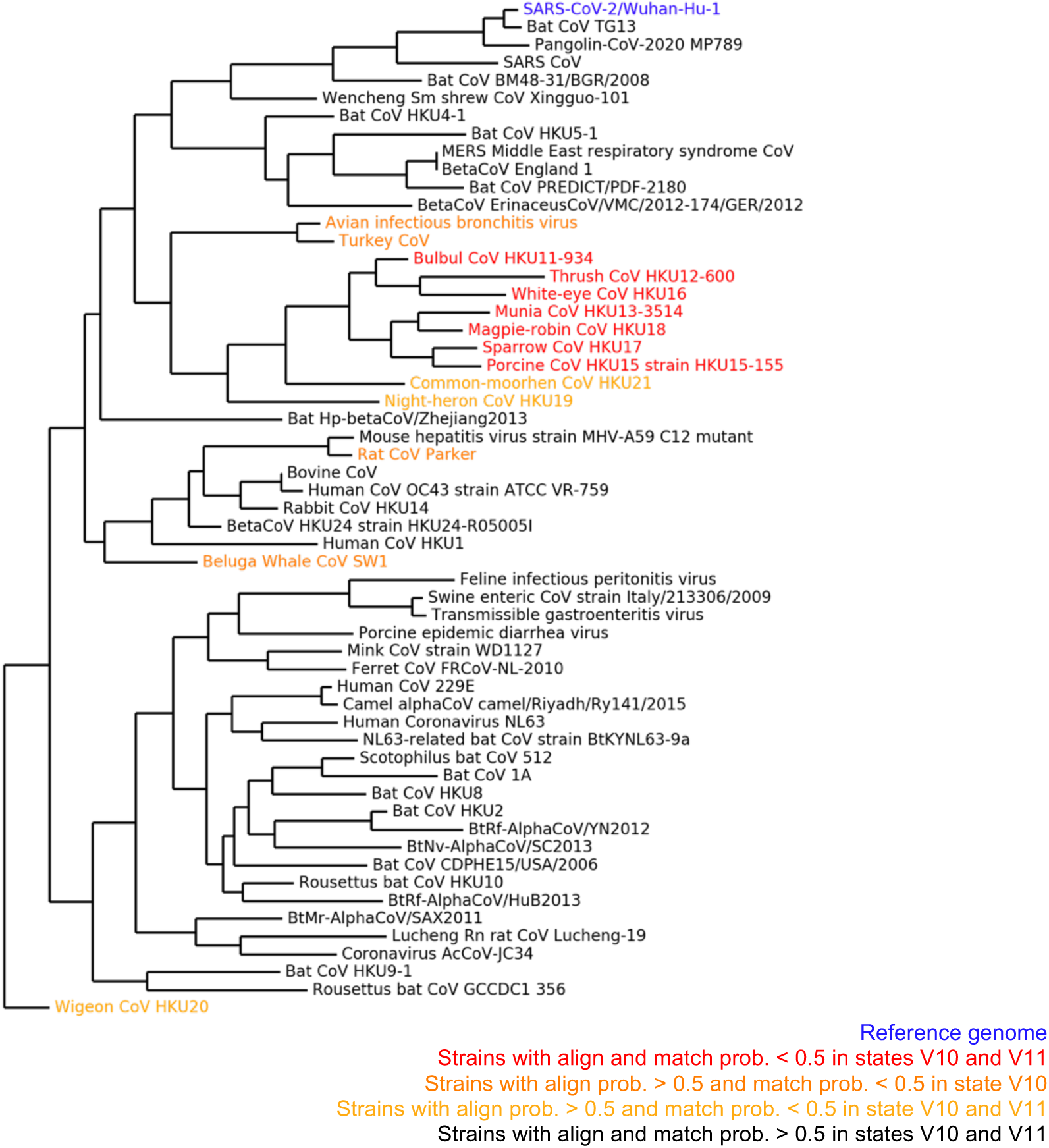
Vertebrate CoV associated with states V10 and V11 in the phylogenetic tree of the vertebrate CoV alignment. Similar to **Fig. 3c** except strains colored according to their align and match probabilities in states V10 and V11. The strain colored in blue is the reference SARS-CoV-2 strain of the alignment, Wuhan-Hu-1. The strains colored in red are those with both align and match probabilities above 0.5 for both states V10 and V11, which include six CoV from avian hosts and a CoV from pig. The strains colored in orange are those with align probabilities above 0.5 and match probabilities below 0.5 for state V10. The strains colored in yellow are those with align probabilities above 0.5 and match probabilities below 0.5 for state V11. The remaining strains in black are those with align and match probabilities above 0.5 for both states.

